# Pushing the limits of structure prediction in regions of disorder using ColabFold: Progress and insights

**DOI:** 10.1101/2024.12.31.630938

**Authors:** Monica D. Rieth

## Abstract

Understanding the impact of amino acid substitutions on protein structure is critical for unraveling mechanisms of protein function and disease. ColabFold, a user-friendly implementation of AlphaFold2, offers an efficient and accessible platform for high-accuracy protein structure predictions. In this study, we explore the utility of ColabFold in predicting structural changes arising from specific amino acid substitutions. Using a structurally well-characterized model protein, mutations were selectively introduced and resulting structural deviations compared to the wild-type conformations. By employing a confidence score embedded in the program called the predicted local distance difference test (pLDDT), which is used to assess a model’s accuracy, together with what is known about the protein’s structure, we identify mutation-induced alterations and determine their potential implications on structural integrity. The results reported here demonstrate that ColabFold is capable of capturing subtle structural perturbations, offering insights into structural stability and the propensity for specific amino acid residues to induce or disrupt secondary structure conformations. We anticipate these studies and others like it will pave the way for high-throughput mutation screening and provide a valuable tool for protein engineering along with hypothesis-driven experimental studies to facilitate our understanding of the behavior of disease-associated mutations.

## INTRODUCTION

Structural biology is the study of molecular structure. It aims to investigate the three-dimensional structures of biological molecules and the relationships between structure and function encompassing a diverse range of experimental techniques, including X-ray crystallography, nuclear magnetic resonance (NMR) spectroscopy, cryo-electron microscopy (cryo-EM), and computational modeling. By revealing the intricate architecture of molecules such as proteins, nucleic acids, and associated complexes, structural biology provides crucial insights into their mechanisms, interactions, and roles in cellular processes.

Cryo-EM has emerged as one of the most powerful tools for studying the structures of viruses, membrane proteins and other challenging biological systems. To complement benchtop techniques, computational modeling and simulations also play a vital role in structural biology. They provide detailed atomic-level models enabling predictions in protein folding, simulating molecular dynamics, and exploring protein-ligand interactions. These methods rely on complex algorithms and powerful computational resources to simulate and analyze the behavior of biomolecules, also enabling a thorough understanding of their structure-function relationships. Understanding the structures of biological molecules has profound implications in drug discovery, as it enables the rational design of therapeutic agents that are aimed at specific molecular targets. Additionally, structural biology contributes to our understanding of diseases, as it provides insights into the molecular basis of disorders, such as cancer, neurodegenerative diseases, and genetic disorders.

Disordered regions of proteins are intriguing and essential components that challenge the traditional view of protein structure. While many proteins fold into well-defined three-dimensional structures, disordered regions, independent of other interactions, lack a kinetically or thermodynamically stable structural conformation, exhibiting a stronger propensity for conformational flexibility and greater degrees of freedom. Despite our more limited understanding of intrinsically disordered protein regions (IDPRs), they play diverse and vital roles in cellular processes, and we are beginning to understand their importance in biological processes. They are involved in protein-protein interactions, signaling pathways, and regulatory functions^1,2^. Short linear interacting motifs (SLiMs) have been identified as regions of heightened interaction for molecular recognition, formation of protein complexes, potential sites for targeted antiviral compounds ^3^. IDRs are also tied to phase behavior phenomena observed both *in vitro* and *in vivo* ^4–8^.

Some of the most debilitating diseases are rooted in the tendencies of some disordered regions to aggregate. IDRs drive aggregation leading to uncontrolled, irreversible fibrillar formation. Formation of protein fibrils underlies the complications that lead to several neurodegenerative diseases such as Alzheimer’s and Parkinson’s ^9–12^. Even though progress continues to be made uncovering the molecular attributes of protein aggregation, our collective understanding of what subsequently leads these complexes to irreversibly form higher order fibrillar structures remains poorly understood. While we can reasonably begin formulating a guiding set of principles governing general properties of intrinsic disorder on a molecular level, questions remain about conditions that mediate intermolecular interactions that drive high-order complex formation.

In disordered protein regions, the amino acid composition is comprised primarily of charged and polar residues. This results in a propensity to interact with the surrounding aqueous environment. Most disordered regions vary in length from short sequences of 5-10 residues, up to much longer sequences of 30 residues or more. Charge distribution along the polypeptide chain is such that the chain resists any tendency to fold onto or interact with itself, lending to a preferentially elongated, unstructured conformation. IDPs that are aggregation-prone have a higher proportion of hydrophobic residues leading to changes in phase behavior^13^. In this same study, authors uncovered features of prion-like domains that promote phase behavior in an attempt to uncover naturally occurring signatures of protein sequences that promote aggregation. Tyrosine and phenylalanine, act as molecular “stickers” and primary drivers of aggregating phase behavior observed in sequences rich in these residues - attributing all other non-aromatic residues as generic spacers that are neutral players in the coupling behaviors that lead to phase separation. The identity and spacing of residues play a key role in intramolecular polypeptide conformation, dictating compaction of a single polypeptide chain. These findings will enable us to better predict which naturally occurring IDRs are susceptible to aggregation and ultimately, fibril formation that leads to cellular neurological disruption. We aim to take this another step further, toward building a set of principles, by probing the primary sequence of a naturally occurring disordered polypeptide to uncover the intramolecular basis of disorder.

In this study, ColabFold was used to investigate residues of the intrinsically disordered C-terminal tail of the human adenosine A2a receptor (hA2a), a GPCR, to predict drivers of secondary structure formation. ColabFold is a readily accessible (requiring minimal computing resources and programming expertise) version of the AlphaFold2 (AF2) program that utilizes the MMseqs2 server to compare protein sequences versus the extensive database network originally employed in earlier versions of the program. AF2 is an AI tool developed by Google DeepMind, that has increasingly grown in popularity as an invaluable resource among structural biologists for predicting protein structures with remarkable accuracy. It utilizes deep learning techniques to predict the two- and three-dimensional structure of a protein from a simple input of its amino acid sequence. AF2’s deep learning architecture is trained on vast amounts of protein structure and sequence data to generate highly accurate structure predictions that have consistently out-performed the competition based on CASP standards^14^. The program employs a two-step process, with the first step involving the generation of a representation of a protein’s amino acid sequence, and the second step employing an attention-based neural network to predict the distances between pairs of amino acids. By combining these distance predictions with optimization algorithms, AF2 produces highly reliable models of protein structures. ColabFold enables a more rapid analysis of protein input sequences by streamlining, through similarity clustering, the reference databases search. It offers comparable, if not better, accuracy in its model predictions and has been quickly adopted by the computational structural biology community. These prediction tools have revolutionized the field of computational structural biology by significantly improving the accuracy and speed of protein structure predictions and guiding experimental design. They have the potential to greatly impact our understanding of protein function, disease mechanisms, and drug discovery, providing invaluable insights into the molecular basis of biological systems. By facilitating experimental design and testing, the once arduous process of protein structure and function elucidation has become increasingly streamlined enabling more effective use of resources.

Together with ColabFold and employing a systematic approach, regions spanning various lengths of the 101-residue cytosolic disordered tail of hA2a were mutated to amino acid residues with different chemical properties to probe their effect, if any, on initiating secondary structure formation. Of the twenty naturally occurring amino acid residues, several were found to be promoters or disruptors of secondary structure formation (α-helical, β-sheet). We chose to use the disordered C-terminal domain and adjacent small 8^th^ helix of hA2a for our analysis because its structure has already been well-characterized experimentally. Therefore, subsequent changes in structure of the variants generated can be critically interpreted with respect to the native sequence. We also defer to previous studies reported in the literature to help guide our query toward relevant questions surrounding the structural basis of disorder. In that way, the results generated from our query could be linked to previously reported experimental studies, lending support to our findings. For example, proline has long been reported to disrupt secondary α-helical structure in proteins. It was found to have a similar effect on the experimental polypeptide sequences tested here. Leucine and alanine promote α-helical structure. These residues are also found enriched in α-helix structures^15,16^. Interestingly, threonine, is suggested to promote β-sheet structure, the only residue found to do so. Using single-chain polypeptides generated from all 20 amino acids, general trends in secondary structure formation were probed and are summarized here. These can be used to help further our understanding of the complexities in protein structure formation and stabilization.

Lastly, IDRs, having no real discernable structure that can be predicted, inherently carry extremely low confidence in their predicted structures. In fact, random coiled and most unstructured regions of even structured proteins, return a very low confidence prediction with ColabFold. Despite these low confidence predictions these programs are enriched for IDRs, which can be informative in and of itself by identifying interesting targets for subsequent analysis. It is estimated that ColabFold-AF2 predicts regions of disorder as well as or even better than many programs designed specifically for such a purpose^17^.

## EXPERIMENTAL METHODS

The full-length amino acid sequence of hA2a was retrieved from the database of GPCR structures (www.gpcrdb.org). The sequence of the cytosolic tail of hA2a was extracted from a snake plot diagram generated from the primary amino acid sequence in the GPCR database program (Figure 2). The “short” (excluding the small predicted interfacial 8^th^ α-helix) intrinsically disordered sequence of hA2a is 101 residues in length (residues 312-412). All mutations were generated manually and their sequence inputted into ColabFold. The “long” sequence included helix 8 (residues 289-412).

ColabFold is an open-source implementation of AlphaFold designed to simplify and accelerate protein structure prediction. ColabFold integrates AlphaFold’s advanced algorithms with optimized features like faster MSA generation using MMseqs2 and support for complex structures, making it accessible to those with limited computational resources and training. *MMseqs2* is a software suite for sequence-sequence and sequence-profile searching and clustering^18,19^. ColabFold-AF2 can achieve high accuracy and has quickly become widely accepted by the computational structural biology community as it has revolutionized structural bioinformatics. Predictions generated in ColabFold-AF2 are assigned a pLDDT value termed the predicted local distance difference test based on the Cα atoms. Regions with pLDDT > 90 are high accuracy predictions; regions with 70 < pLDDT < 90 are confident predictions (with respect to their backbone orientation); regions with 50 < pLDDT < 70 should be interpreted with caution (low confidence); and regions with pLDDT < 50 are generally not considered to be accurate predictions with very low confidence. The structure of these regions should not be interpreted. Disordered protein regions typically fall below pLDDT < 50. A working copy of the ColabFold notebook can be copied from https://alphafold.colabfold.com to Google Drive. Alternatively, ColabFold can be operated directly from the open source github notebook (https://github.com/sokrypton/ColabFold) ^20^. Results generated from query sequences were downloaded and saved locally for analysis.

The notebook setup for each prediction was loosely based on a recent protocol that can be found at https://doi.org/10.21203/rs.3.pex-2490/v1 ^21^. It describes how to set up a modeling experiment, explains the parameters, and output data. In these experiments, a simple monomer prediction was utilized to assess polypeptide secondary structure of carefully selected query sequences aimed at addressing fundamental questions about primary amino acid sequence structure and propensity to promote structural disorder. The ColabFold notebook predicts the top five models. By default, relaxation was left enabled for the final structure predictions. Models were assessed using the pLDDT confidence metric. These predictions have been found to be useful in informing regions of disorder relatively well^17^. This parameter was used to assess local changes in the predicted structures of query (experimental) sequences. Wherever possible and relevant, changes in the certainty of structural predictions were assessed by examining pLDDT values of variants compared to the WT. Other parameters such as the predicted aligned error (PAE) can help inform interdomain positioning in a predicted structure. In this case, since we are working with primarily disordered single polypeptide domains, this parameter provides little added meaning to the interpretation of our results. Sequence coverage for each entry was also considered in evaluating each of the results. In most cases, the default settings were left unchanged, and sequence coverage was adequate. A minimum of 30 sequences is recommended for accurate predictions to be made. This value was met or exceeded for most of the query sequences in this analysis. A schematic overview in Figure 1 shows a general summary of the workflow followed in our study.

**Figure 1.**
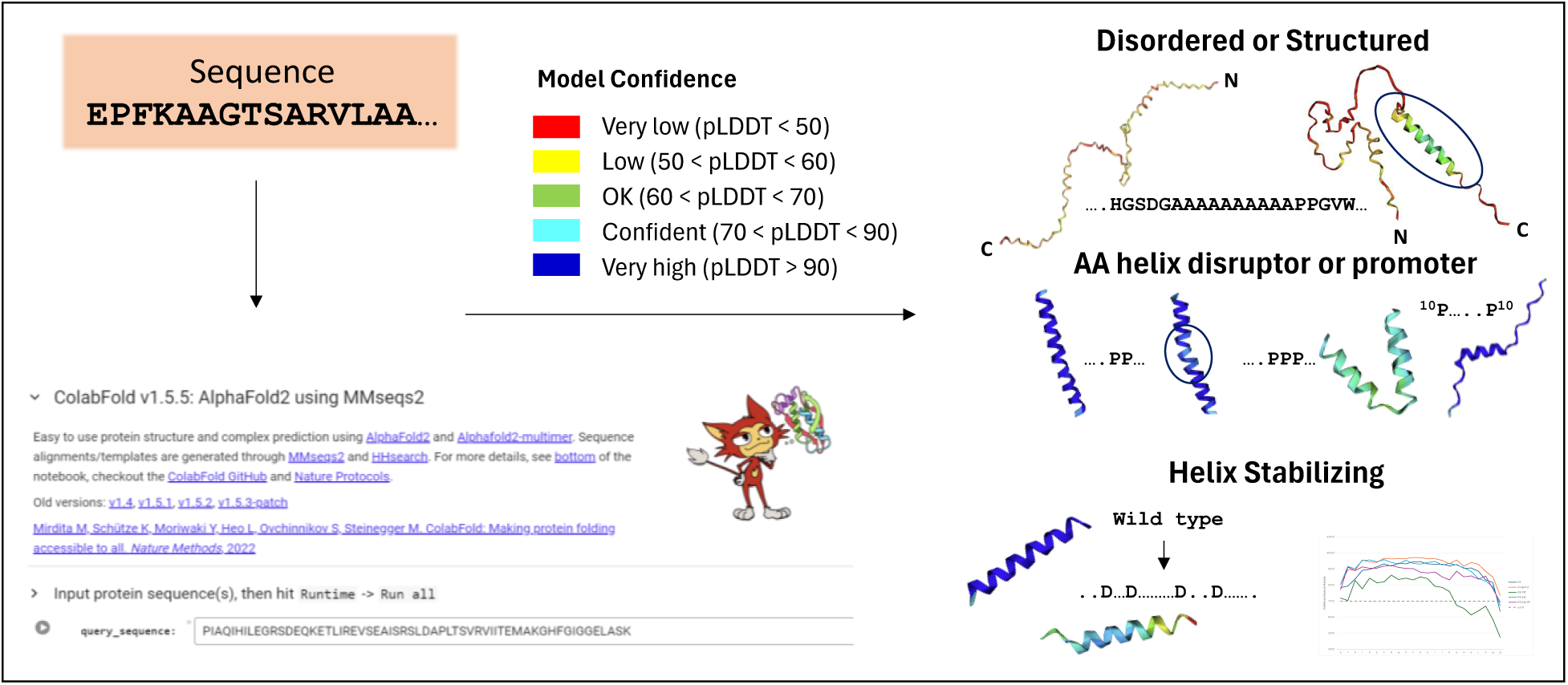
Overview of ColabFold workflow. Query sequences are inputted into the ColabFold notebook. Models are assessed for confidence using the color-coded scale, which maps to the 3D molecular view generated by the program. Regions predicted with low confidence are colored in red-yellow (pLDDT < 60). These regions are not likely to be structured (disordered). Models in blue carry high confidence (pLDDT > 90) in their predictions and can be reasonably interpreted as well as models in light blue-green (pLDDT > 60). Specific amino acid residues were tested in query sequences to assess their effect on known structured and disordered regions. Variants of the C-terminal disordered tail of the human A2a receptor were assessed by evaluating the 3D structural model and analyzing both global and local structural changes resulting from sequence substitutions. Other residues were introduced to assess their respective changes to the structured helix 8 (also native to hA2a). Changes were made to probe the effects of charged residue substitutions in helix 8. pLDDT values were analyzed for variants in helix 8 to determine which substitutions stabilized or destabilized alpha helical structure.

Query sequences were designed using the wild-type disordered C-terminal domain of hA2a and the small 8^th^ helix. The primary sequence of each mutant was inputted individually into the ColabFold notebook. pLDDT scores for each residue were evaluated comparatively for each mutant to determine relative change in confidence values of the predicted structure. An increase in the confidence value of a residue at a substituted site compared to the value of the wild-type residue was determined. The change in confidence value of each substituted residue corresponding to the predicted structure (for the mutant) indicates a positive or negative structural effect. This was carried out for helix 8, only, where initial predictions to the wild-type sequence could be made with very high confidence. Generally, negative changes are attributed to destabilizing effects or increased uncertainty in the model resulting from changes in the variant primary sequence. For the C-terminal disordered region and its variants, where predictions already carry very low confidence values, the 3D model rendering was evaluated for changes in structure and associated confidence values using the corresponding color coding already integrated into the program. This was preserved as the color coding is uniquely meaningful to the ColabFold program and helps to inform our analysis of the resulting variant effects on structural disorder.

The data were exported from the ColabFold program along with the color-coded structural renderings. pLDDT values for files corresponding to helix 8 variants were exported in .txt format and opened in MS Excel where they were plotted for each residue for comparison with the WT sequence (x-axis). Values on the y-axis correspond to confidence values between 40 and 100. Values less than 50 are considered very low confidence and caution should be used to make any meaningful interpretation about structure. pLDDT values for each residue in all variants of helix 8 were above this threshold. An arbitrary threshold of 70 was indicated by a dotted gray line. Values above 70 are an indication of confidence in structure. Values for each residue of the disordered C-terminal domain lie below this threshold making structural predictions of its variants too ambiguous to decipher with any reasonable level of confidence using pLDDT values. Therefore, we defer to the structural model produced to determine if any appreciable secondary structure could be detected in the variants tested.

## RESULTS AND DISCUSSION

The structures of all query sequences were predicted using ColabFold^20^. Detailed instructions on its use are described in the notebook. The test “query” sequences were inputted into the “query sequence” prompt space in the notebook and all cells were executed simultaneously: “Runtime” → “Run all.” Default settings were unchanged. Generally, more iterations improve the quality of the final prediction. However, for our test sequences, there was no obvious benefit to increasing the number of iterations. Relaxation parameters were left on. Results for the top five ranked models were automatically downloaded in a zip file that includes the pdb file of the predicted structural model along with color-coded confidence rankings. Confidence values (pLDDT) equal to or above 70% were considered significant predictors of structure formation and further probed. The top ranked models are presented herein. Sequences showing little to no secondary structure are strong indicators of intrinsic disorder. They resemble more “spaghetti-like” topology with extremely low confidence rankings, often equal to or below 50%. The C-terminal domain of the human adenosine A2a receptor, a G-protein coupled receptor (GPCR), is a cytoplasmic facing region of the protein that has been shown experimentally using solution NMR to be disordered in the absence of protein-protein interactions^22^. This region was used as the basis to initiate analysis of secondary structure-inducing amino acid residues.

### Benchmarking amino acid residues

All twenty amino acids were tested using poly-AA sequences to determine which residues had a higher propensity to promote secondary structure formation, by themselves, “promoters.” As well as which residues disrupt secondary structure formation, “disruptors.” From here on we incorporate the naming convention, “order-promoting” and “disorder-promoting” to describe amino acids with the respective effects^23^. A 25-mer poly-AA sequence was generated for each amino acid. Results are summarized in Supplementary Tables S1-3. Poly-AA sequences that were predicted to have secondary structure were classified as promoters or order-promoting while those that exhibited ambiguous or neutral structural features were classified as disruptors or disorder-promoting. Consequently, incorporation of these sequences into the intrinsically disordered region of A2a were expected to result in the corresponding effect on the localized structure. Overall, all hydrophobic residues: Leu, Iso, Ala, Val, Phe, Trp, Tyr and Met were promoters of α-helical secondary structure (Supplemental Table S3). Glu, Lys, Arg, and Gln (Supplementary Tables S1,S2) also adopt helical structure. The remaining amino acids, Ser, Asn, Asp, Gly, Pro, His showed little propensity to adopt secondary structure. Cysteine appears to be more “ambivalent.” Interestingly, Thr, was the only amino acid to exhibit β sheet structure.

ColabFold-AF2 has not been trained or validated to predict the effects of point mutations on resulting protein structure. Therefore, the destabilizing or stabilizing effects of single amino acid substitutions on protein structure cannot be accurately predicted. The program is also unable to predict thermodynamic changes, *ΔΔG*, in comparative protein structures. AlphaFold claims, “for regions that are intrinsically disordered or unstructured, AlphaFold is expected to produce a low-confidence model structure prediction (pLDDT < 50). AlphaFold is useful in identifying such regions, but the program does not make any claims about the relative likelihood of different conformations” (cited from: https://alphafold.ebi.ac.uk/faq).

Disordered regions are known to be extremely dynamic in nature, assuming many possible conformations. Because disordered regions already produce very low confidence structure predictions, rigorous analysis of pLDDT values between predictions of variants and WT sequences provides little insight. We can, however, qualitatively inspect the resulting models for localized and global changes in structure irrespective of their relative confidence values. For known structured regions, e.g. hA2a helix 8, we can better analyze changes in pLDDT as a function of variations in the primary sequence because these are already predicted with high confidence.

### Design of query sequences

Of the twenty amino acids, Leu, Ala, Gly, and Pro were chosen for further analysis due to their well-documented induced effects on protein structure. Alanine and leucine are known promoters of α-helical secondary structure^15,24–26^. Synthetic polypeptide sequences comprised almost entirely of leucine and alanine residues, called “WALPs”, adopt^24,27^ α-helical structures of various lengths depending on the number of residues. These residues were chosen for subsequent analyses because of their well-characterized effects on structure. They were tested in ColabFold by introducing polyalanine and polyleucine substitutions along the length of the disordered C-terminal domain. Proline and glycine are known helix disruptors. This is also consistent with the reported findings in Table S2 where both appear to adopt an extended disordered polypeptide chain. While point mutations are said to be beyond the capability of ColabFold predictions, we report one case where the introduction of just one proline residue into hA2a helix 8 was sufficient to alter the structure (see below). In general, however, single point mutations in query sequences were avoided for the reasons mentioned above. Instead, our analysis focuses on investigating the effects of poly-AA sequences substituted into the C-terminal disordered domain and helix 8.

Poly Q sequences are a genetic anomaly that occur resulting in the production of poly Q sequences into native protein structures. The introduction of these sequences has been linked to the abnormal protein behavior underlying a number of neurodegenerative diseases ^28,29^. The structural effects of poly Q incorporation were tested in helix 8 to verify if poly Q could induce disorder.

Lastly, disordered proteins are stabilized by the presence of highly charged amino acid residues. The distribution of native sequence charges and their relative charge densities prevents them from adopting close conformations required for stabilized secondary structures. This phenomenon continues to be an ongoing area of investigation so that we can better predict protein sequences that are most susceptible to adopting disorder and subsequently aggregating into high-order structures - which lies at the heart of many diseases. We began probing the question - how altering the charged residues of helix 8 effects its structure - by analyzing model structures produced by ColabFold and comparing relative pLDDT values with the wild-type (WT). This tells us if the variants tested lead to more or less stabilized structures using associated confidence values of the variant models. Lower confidence values indicate a possible destabilizing effect and increased uncertainty in structure while higher confidence values indicate a possible stabilizing effect. These changes enable insights only, no thermodynamic data for the structures was determined, as ColabFold does not produce *ΔG* values or any other thermodynamic properties of its models.

### Prediction of the native C-terminal domain of adenosine A2a

The disordered region of the C-terminal domain of adenosine A2a encompasses approximately the last 100 residues of the protein beginning at position 312 through 412. Figure 2 shows the ColabFold predicted structure of the top five ranked models for full-length hA2a. The fully disordered region is boxed to illustrate the region of interest in the following analysis. Not surprisingly, based on the predictions, the signature seven transmembrane helices of hA2a are shown to be very well-structured (dark blue-left, indicating highest confidence prediction) including helix 8. The region encompassing residues 312-412 is predicted, with lowest probability, to be highly unstructured. These predictions are consistent with the experimental evidence to date^22^.

**Figure 2.**
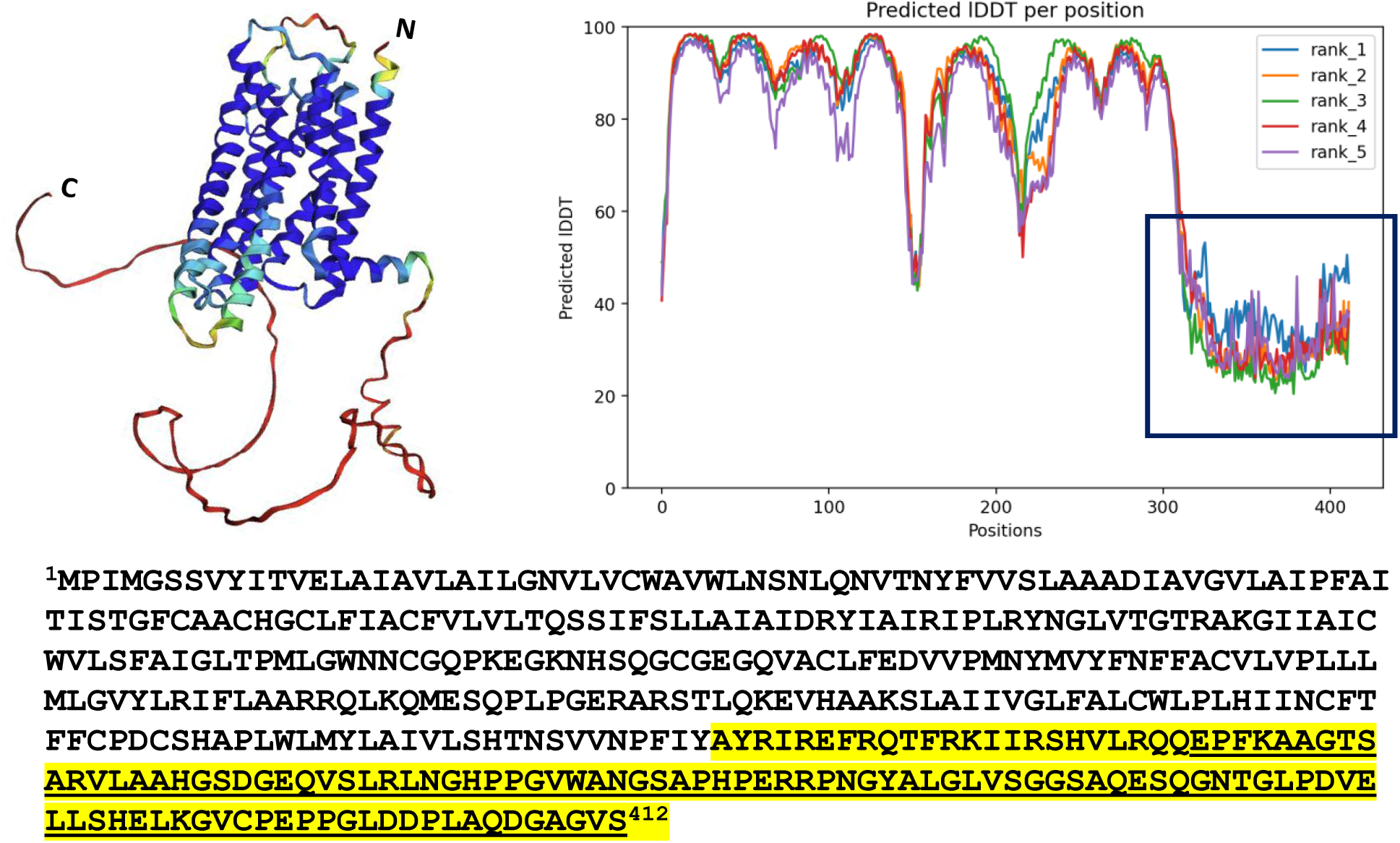
Predicted model and primary sequence of the human A2a receptor. The C-terminal domain, short, (yellow-underlined), residues 301-412, is a low confidence prediction, < 60%, because it is disordered. ColabFold predicts disordered regions with very low confidence because they do not assume any appreciable structure and cannot be modeled well by the program. The intra- and extra-cellular loop regions are also relatively disordered and present with lower confidence, < 80%, compared to the transmembrane domains, > 90%. The transmembrane regions are well structured and depicted with very high confidence (in blue-left), >90%. The pLDDT values for the top five model structures produced are overlayed (right).

### Polyalanine and polyleucine scanning of C-terminal domain

To begin this analysis sequence constructs of the C-terminal domain both with and without the cytoplasmic 8^th^ helix were analyzed. For each construct 10 query sequences were created where 10 alanine residues were substituted along the length of the disordered region of each construct in a consecutive stepwise manner with no overlap. Sequences are summarized in Table 1.

**Table 1.**
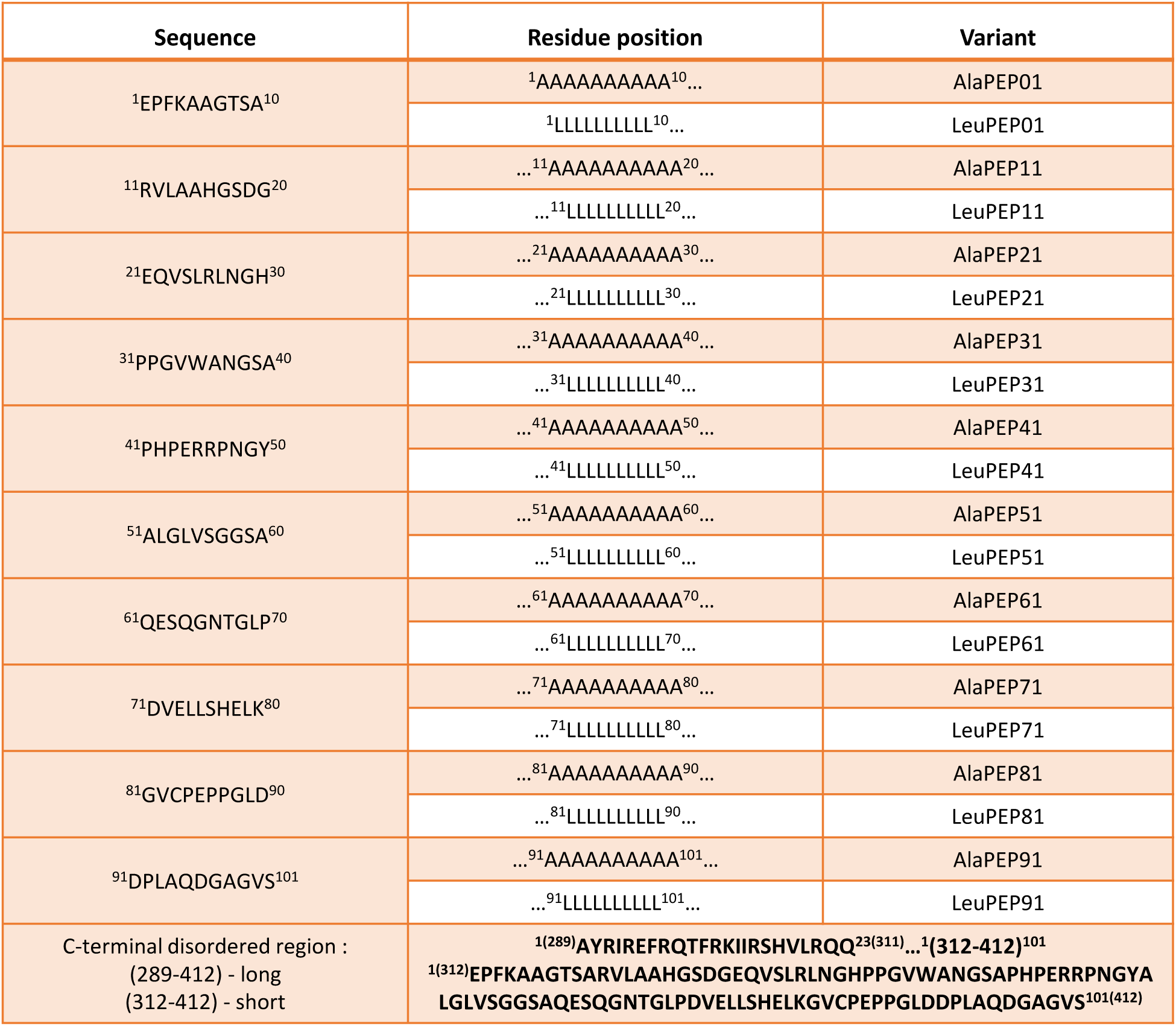
Polyalanine and polyleucine query sequences.

| Sequence | Residue position | Variant |
| --- | --- | --- |
| <sup>1</sup>EPFKAAGTSA<sup>10</sup> | <sup>1</sup>AAAAAAAAAA<sup>10</sup> ... | AlaPEP01 |
|  | <sup>1</sup>LLLLLLLLLL<sup>10</sup> ... | LeuPEP01 |
| <sup>11</sup>RVLAAHGSDG<sup>20</sup> | ...<sup>11</sup>AAAAAAAAAA<sup>20</sup> ... | AlaPEP11 |
|  | ...<sup>11</sup>LLLLLLLLLL<sup>20</sup> ... | LeuPEP11 |
| <sup>21</sup>EQVSLRLNGH<sup>30</sup> | ...<sup>21</sup>AAAAAAAAAA<sup>30</sup> ... | AlaPEP21 |
|  | ...<sup>21</sup>LLLLLLLLLL<sup>30</sup> ... | LeuPEP21 |
| <sup>31</sup>PPGVWANGSA<sup>40</sup> | ...<sup>31</sup>AAAAAAAAAA<sup>40</sup> ... | AlaPEP31 |
|  | ...<sup>31</sup>LLLLLLLLLL<sup>40</sup> ... | LeuPEP31 |
| <sup>41</sup>PHPERRPNGY<sup>50</sup> | ...<sup>41</sup>AAAAAAAAAA<sup>50</sup> ... | AlaPEP41 |
|  | ...<sup>41</sup>LLLLLLLLLL<sup>50</sup> ... | LeuPEP41 |
| <sup>51</sup>ALGLVSGGSA<sup>60</sup> | ...<sup>51</sup>AAAAAAAAAA<sup>60</sup> ... | AlaPEP51 |
|  | ...<sup>51</sup>LLLLLLLLLL<sup>60</sup> ... | LeuPEP51 |
| <sup>61</sup>QESQGNTGLP<sup>70</sup> | ...<sup>61</sup>AAAAAAAAAA<sup>70</sup> ... | AlaPEP61 |
|  | ...<sup>61</sup>LLLLLLLLLL<sup>70</sup> ... | LeuPEP61 |
| <sup>71</sup>DVELLSHELK<sup>80</sup> | ...<sup>71</sup>AAAAAAAAAA<sup>80</sup> ... | AlaPEP71 |
|  | ...<sup>71</sup>LLLLLLLLLL<sup>80</sup> ... | LeuPEP71 |
| <sup>81</sup>GVCPEPPGLD<sup>90</sup> | ...<sup>81</sup>AAAAAAAAAA<sup>90</sup> ... | AlaPEP81 |
|  | ...<sup>81</sup>LLLLLLLLLL<sup>90</sup> ... | LeuPEP81 |
| <sup>91</sup>DPLAQDGAGVS<sup>101</sup> | ...<sup>91</sup>AAAAAAAAAA<sup>101</sup> ... | AlaPEP91 |
|  | ...<sup>91</sup>LLLLLLLLLL<sup>101</sup> ... | LeuPEP91 |
| C-terminal disordered region :<br>(289-412) - long<br>(312-412) - short | <sup>1(289)</sup>AYRIREFRQTFRKIIRSHVLRQQ<sup>23(311)</sup>...<sup>1(312-412)</sup><sup>101</sup><br><sup>1(312)</sup>EPFKAAGTSARVLAAHGSDGEQVSLRLNGHPPGVWANGSAPHPERRPNGYA<br>LGLVSGGSAQESQGNTGLPDVELLSHELKGVCPEPPGLDDPLAQDGAGVS<sup>101(412)</sup> |  |

Three of the 10 variants for the long and short hA2a sequences give some indication of secondary structure adoption with the substitution of polyalanine sequences. The structural models are shown in Fig. 3. There does not appear to be any difference in the induced effects for the long, panels D-F, or the short, panels A-C, constructs. Due to the highly disordered nature of the C-terminal domain, confidence values fell below 60. A comparison of pLDDT values between the WT and the variants provided little added meaning because both confidence predictions lie below the threshold cutoff recommended for interpretation (pLDDT < 60). This was also true of polyleucine variants.

**Figure 3.**
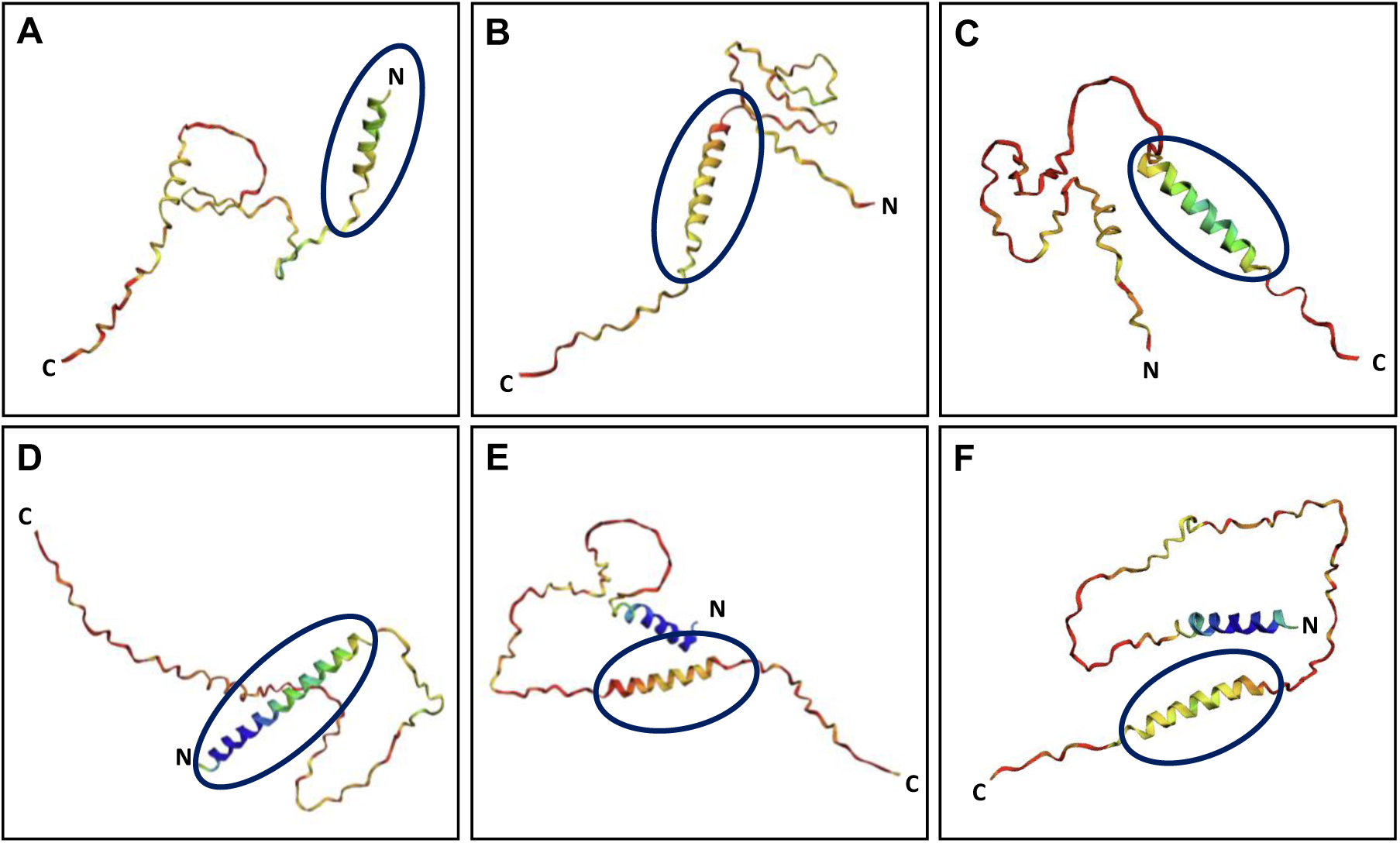
Polyalanine induces localized helical structure. Panels A-C illustrate the short C-terminal domain of hA2a without the 8^th^ helix. Three out of 10 query sequences are shown to adopt alpha helical formation upon introducing tracts of alanine residues: A) AlaPEP01, B) AlaPEP61, C) AlaPEP81. Panels D-F illustrate the long C-terminal domain of hA2a with helix 8. The same three sequences show α-helical formation upon inclusion of alanine residue tracts: D) AlaPEP01, B) AlaPEP61, C) AlaPEP81.

Leucine residues were introduced to the disordered C-terminal domain. In light of the similarities of both the long and short constructs, substitutions were made only to the long construct going forward in our analyses. In all 10 query sequences, polyleucine substitutions promoted helix formation in regions where it was introduced to the otherwise disordered C-terminal domain. Our benchmark analysis showed leucine is sufficient, by itself, to assume a single α-helical structure (Table S3). Further, incorporation of alanine residues to produce a hybrid “LA” sequence synonymous to the reported “WALP” synthetic structure shows good agreement in ColabFold predictions to strongly assume α-helical structure as well^32^ (not shown). To produce these hybrid constructs, a 25-residue polyleucine query sequence was inputted into ColabFold followed by several other permutations of hybrid sequences: a 53-residue polyalaleucine sequence flanked on either end by 15 alanine residues. Additional sequences of varying lengths were queried and produced similar results (not shown). All were predicted to have α-helical secondary structure with very high confidence based on pLDDT scores. The minimum number of residues reportedly required to assume single-chain α-helical secondary structure containing a single turn is 4-5^33^. In nature, however, the average length of α-helical regions is found to be closer to 11 residues^34^. The minimum requirement for a polyleucine α-helix was approximately 8 residues for a two-turn helix in ColabFold (not shown).

*In vivo* and *in vitro*, polyleucine α-helices and their variants have been studied as model transmembrane peptides to understand how membrane proteins interact with one another in a lipid bilayer environment synonymous to the plasma membrane. Due to their hydrophobic nature, they readily partition into membrane lipid bilayers^24,27^. On the contrary, a defining feature of IDPs is their high net charge at physiological pH due to an abundance of charged residues. We hypothesize that introducing more hydrophobic residues to a disordered protein would promote structure formation. In all cases, this turned out to be true. Localized helical structure was adopted at sites where polyleucine sequences were substituted (Fig. 4). These findings are consistent with previous reports looking at the propensity of each residue to participate in structured or unstructured proteins^23^.

**Figure 4.**
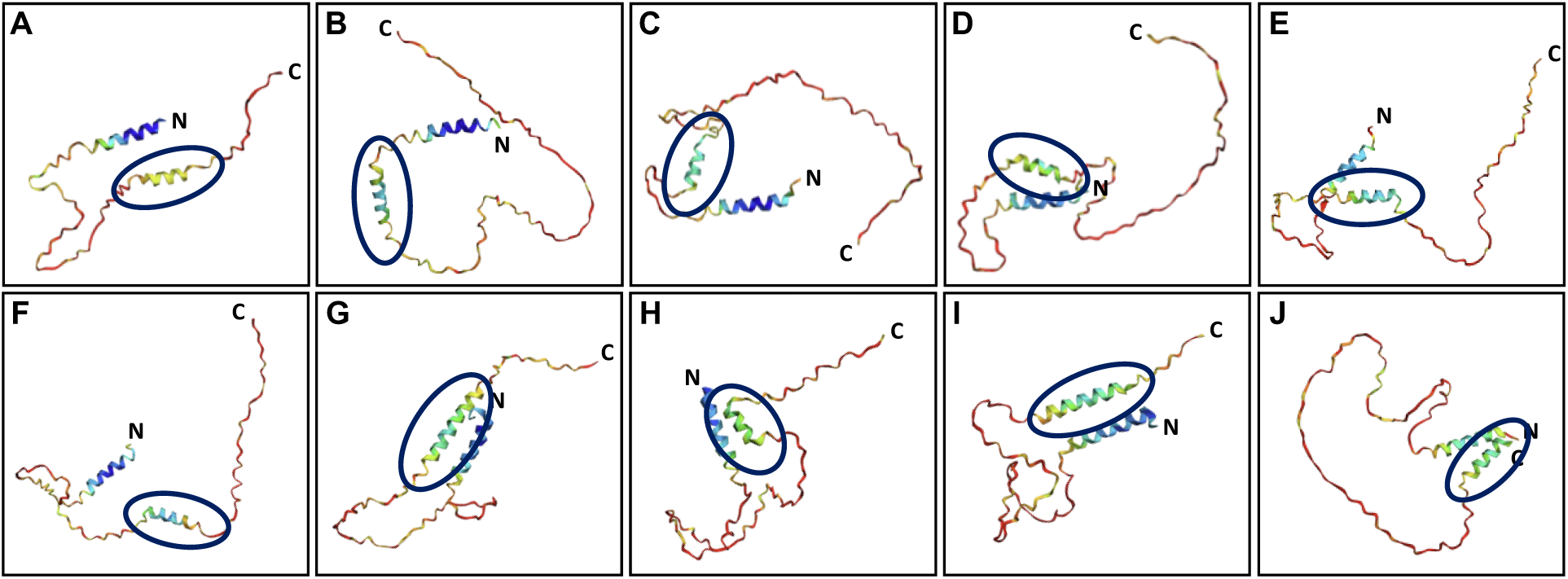
Leucine is a strong promoter of helical structure. Polyleucine sequences were substituted at sites in the same manner as alanine (refer to Table 1) along both the long and short C-terminal domains of hA2a. In all cases, helical structure was detected at sites of substitution. Here, the long C-terminal region for all 10 query sequences is shown. Synonymous sites in the short C-terminal domain yield similar results (not shown).

### Proline is disorder promoting

It is well established that proline is a unique amino acid with distinct structural properties that make it a notable disruptor of α-helices in proteins. Unlike other amino acids, proline’s side chain forms a cyclic structure by covalently bonding to its own amino group, creating a pyrrolidine ring. This rigid, non-rotatable structure imposes a fixed angle on the peptide backbone, significantly limiting its conformational flexibility. In α-helices, hydrogen bonds between the backbone amide groups stabilize the helical structure. Proline’s cyclic structure prevents it from participating in these hydrogen bonds because its amide group lacks a hydrogen atom necessary for bonding. Additionally, the fixed φ angle of proline (∼ -65°) is incompatible with the typical angles found in α-helices, further disrupting the helical arrangement^35^. Consequently, when proline is present in a polypeptide sequence, it often acts as a “helix breaker,” introducing kinks or disruptions in helical regions^27,36^. This disruption can play a functional role in protein structure and dynamics by imparting localized backbone rigidity when present. It can also dramatically affect a protein’s functional behavior^37^. Despite its helix-disrupting properties, proline is essential for creating turns, loops, irregular regions, and its own unique helical structure (polyproline II helix) contributing to the diversity and complexity of protein folding. In intrinsically disordered regions, the significance of proline has been explored. It is disruptive to secondary structure formation by altering backbone chain compaction^38^, and thereby promoting disorder. These findings are consistent with earlier studies that looked at the role of individual amino acids in promoting or disrupting helical activity. Proline was found to be a helix disruptor or “disorder promoting” based on a statistical analysis using multiple data sets comparing the frequency with which all 20 amino acids occur in a sample set of proteins^23^. One study suggests the placement of a proline residue at either the N- or C- terminus of pre-structured recognition motifs (PreSMos or MolRFs), localized sites in disordered proteins with high propensity to assume secondary structure upon binding, induces disorder. MolRFs are short linear motifs that give rise to recognition-binding (“coupled folding-binding”) properties exhibited in IDPs where disorder-to-ordered transitions commonly take place^38^. In many cases, MolRFs determine the function of an IDPR and which interactions it will participate in, protein-protein and protein-DNA interactions^23^. Structurally, many hydrophobic and aromatic residues are highly conserved in these motifs, indicating an evolutionary important functional role.

Using ColabFold, proline residues were introduced to the small 8^th^ helix of hA2a semi-systematically. Since AF2 cannot accurately capture structural perturbations that result from single point mutations, a minimum of two to three consecutive amino acid substitutions were required to produce any detectable changes in the structure of the query sequences. In our case, this was found to be sequence dependent. The addition of a single point mutation, proline, to the hA2a 8^th^ helix resulted in a localized disruption (Supplementary Fig. S3) depending on where the substitution took place. For example, a substitution at position F299P did not result in a detectable “kink” in the predicted 3D helical structure, but it did noticeably reduce confidence in the prediction. On the other hand, a R300P point mutation disrupted the predicted 3D structure, and reduced confidence at the substitution site. Continuous proline-scanning (introducing three proline residues at various points of substitution) of helix 8 showed disruption that coincides with points of substitution. However, the disruption occurred locally while the remaining polypeptide retained its α-helical properties (not shown). These findings are consistent with previous reports of localized proline-induced structural changes as well as our polyproline 25-mer model prediction (Supplementary Table S2).

Until recently, AF2 reportedly failed at predicting the effects of single residue point mutations when investigated in the context of fluorescence experiments^30^. However, a study by McBride and co-workers challenges this notion by showing that a subset of structure pairs differing by 1-3 point mutations reported in the PDB correlates well with perturbations in their AF2 predicted structures^31^. This supports our claim that the effects of point mutations are sequence and position dependent.

The effect of proline was further probed looking at a polyleucine/polyalanine hybrid helix (Leu*_x_*-Ala*_y_*-Pro*_z_*-Ala*_y_*-Leu*_x_*). Again, introduction of a single proline residue showed little to no effect on structure, while two consecutive proline residues gave rise to a “kink” in the helix, and three consecutive prolines resulted in a near complete turn in the helix backbone (Supplementary Fig. S1). Interestingly, introduction of an experimentally-determined three-residue turn comprised of glycine, isoleucine, and proline (GIP) also results in a near complete or U-shaped turn to the Leu-Ala hybrid helix (Supplementary Fig. S2)^39^.

### Poly Q sequences

Polyglutamine (poly Q) diseases are a group of neurodegenerative disorders caused by abnormal expansions of CAG trinucleotide repeats in specific genes, which encode for proteins with an elongated polyglutamine tract ^40–42^. The expanded poly Q tracts lead to protein misfolding, aggregation, and a toxic gain-of-function mechanism. At the molecular level, these misfolded proteins form intracellular aggregates, disrupting cellular functions. Additionally, they trigger cellular stress pathways and neuroinflammation, contributing to progressive neuronal dysfunction and cell death. We investigated the effects of introducing poly Q sequences to helix 8 to determine if there were any obvious destabilizing effects on the predicted structure. Our initial query of the 25-mer sequences of amino acids revealed that polyglutamine (Q) produces a helical structure with extremely high confidence (Fig. 5, panel F). Replacement of all charged residues in helix 8 with Q also produces a helical structure with high confidence (Fig. 5, panel A). To reconcile these findings with previous reports, a series of additional variants was explored. Poly Q sequences comprised of 11-12 mer polypeptides (consecutive glutamine residues) were substituted at three different sites in helix 8, 1) replacing the first 12 residues, Q_12_-WT, 2) replacing the middle 11 residues, WT-Q_11_-WT, and 3) replacing the last 11 residues, WT-Q_11_. In all three poly Q variants we see destabilizing effects (Fig. 5) indicated by reduced confidence levels and greater uncertainty in the prediction. Fig. 5 shows that when a poly Q_11-12_ sequence is introduced toward the N-terminus there is more uncertainty in the helical structural prediction, panels B and D. Our benchmark analysis suggests that polyglutamine (poly Q) is order-promoting. In fact, ColabFold predicts poly Q helical structure with greater certainty than the WT helix 8. This suggests that surrounding residues in regions of poly Q substitution may be critically important in preserving helical structure when poly Q mutations arise.

**Figure 5.**
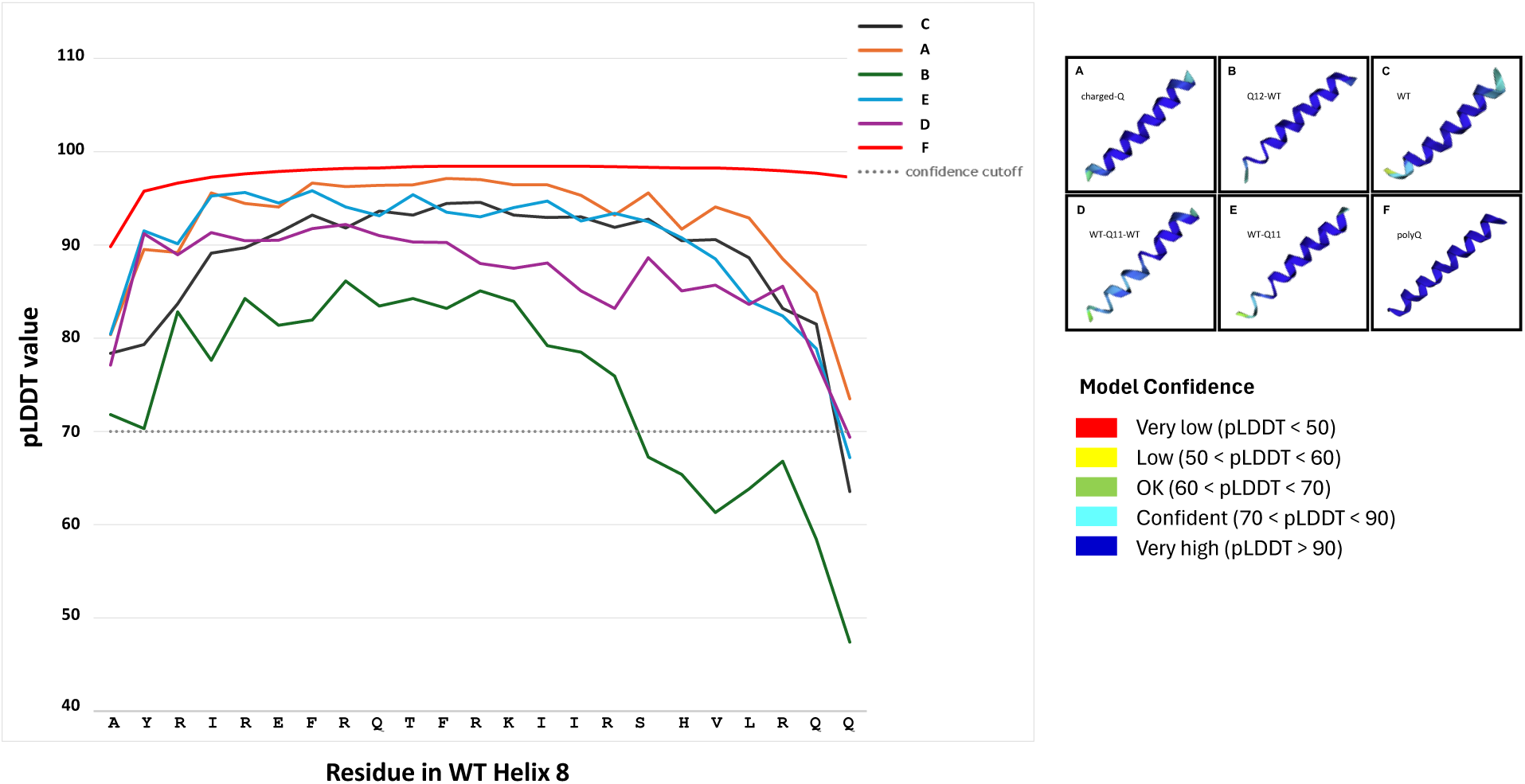
Confidence values per residue for poly-Q variants of Helix 8. The wild-type primary sequence of helix 8 is plotted on the x-axis and associated pLDDT value for each residue in the models is plotted on the y-axis. Leading poly Q sequences (green & purple-left) shows the most significant decrease in confidence value per residue for each variant. Poly Q (panel F) gives the highest confidence for helical structure. Confidence in the predictions gradually increases as the poly Q tracts are placed near the C-terminus of the helix. Whereas structural uncertainty increases as poly Q tracts are placed nearer to the N-terminus. The model confidence color coding corresponds to the variant models in panels A-F only. The model structures in each panel are shown C terminus to N terminus from left to right.

With that in mind, it is likely that certain protein structures are inherently more susceptible than others to alteration by poly Q incorporation based on the nature of their chemical properties. Further deciphering these properties could help identify which proteins are more vulnerable to poly Q mutations that lead to disease-causing aggregation.

### Examining the impact of charged residue substitutions

In studying the molecular composition of IDPs and IDRs, several patterns have begun to emerge. Order promoting residues often include non-polar amino acids and are found buried within the core of globular proteins^23^. Disorder-promoting residues are usually polar and often encompass charged residues. In globular proteins, the latter are typically found at the surface. Protein tertiary structure is driven and stabilized by the hydrophobic residues that lie at a protein’s core. When a polypeptide chain is heavily comprised of charged residues, this prevents the polypeptide from collapsing and folding into a compact structure. Instead, it prefers to remain unstructured and in elongated form giving rise to IDPs. To probe the contribution of charged residues to the disordered C-terminal domain and helix 8, selected substitutions were made to assess the effects on α-helical structure. Variants are summarized in Table 2. Residues were selected to specifically alter the physiochemical properties that would result in complete neutralization of all charged residues, positively charged residues, and negatively charged residues only. Native charged residues were altered to the opposite charges as well, e.g. positive to negative changes and negative to positive changes. Some residues were selected based on their benchmarked order- or disorder-promoting characteristics (Supplementary Tables S1-3). Interestingly, the results observed for the substituted residues tested were consistent with the observed induced structural effects for each respective amino acid. For example, aspartate and glycine are both disorder-promoting based on the findings from our benchmark analysis. Introducing these residues into helix 8 produced increased uncertainty in the prediction. This observation holds true for variants where these residues were substituted for positively charged residues, Fig. 6 – Panel A. It was especially pronounced when aspartate residues were substituted for uncharged residues in Fig. 6 – Panel D where the structure of helix 8 becomes almost completely disordered (Supplementary Fig. S4 – Panel I). Histidine is also disorder-promoting and increasing disorder was observed when more residues were incorporated into the sequence of the helix (Fig. 6 – Panel D). In panel B of the same Fig., only one negatively charged residue was substituted with histidine (negative to positive substitution) resulting in much less significant disruption to the helical structure, however, slight disruption occurs toward the C-terminal end of the helix when compared to the WT.

**Figure 6.**
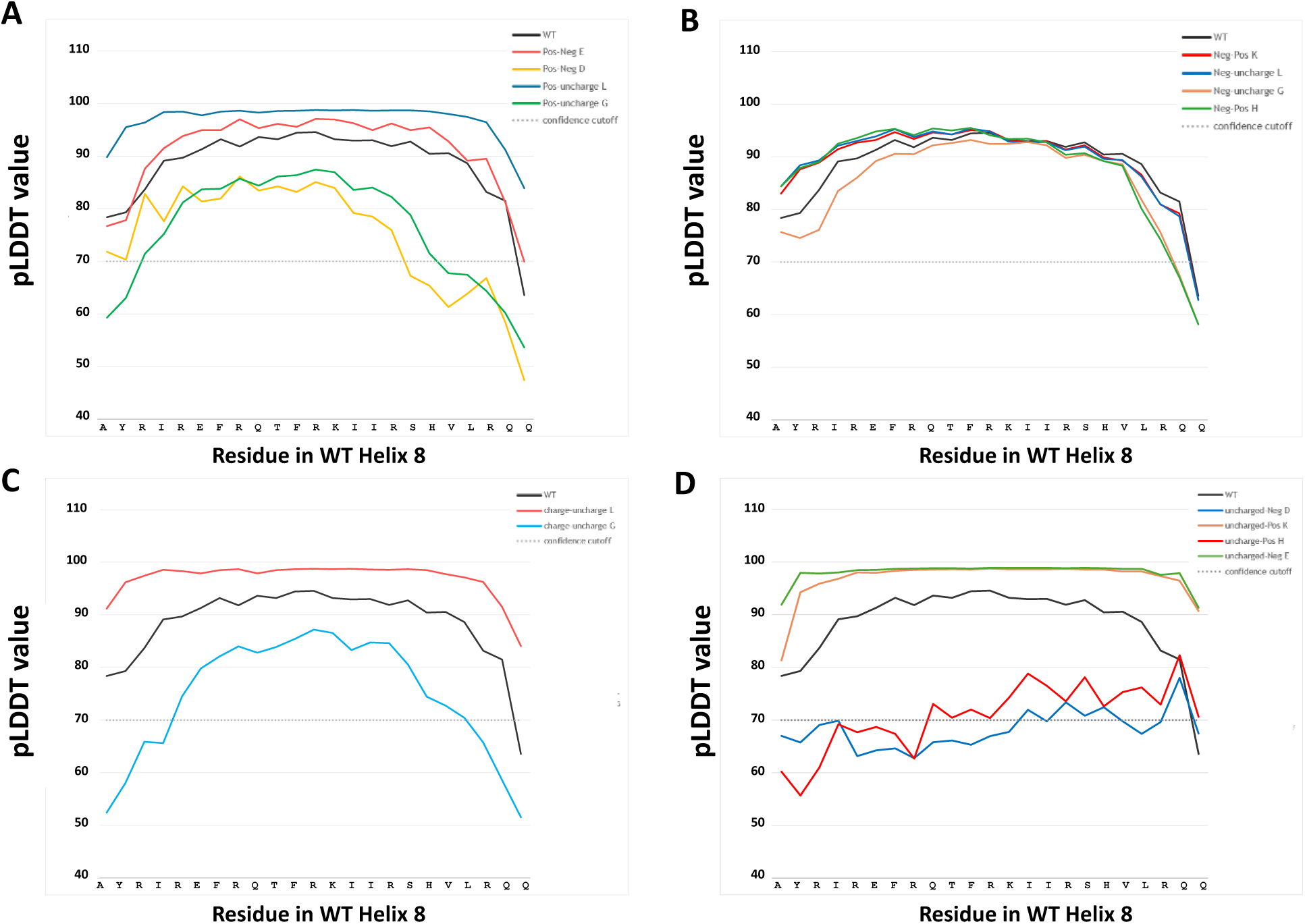
Confidence values per residue for charged residue variants of Helix 8. The wild-type primary sequence of helix 8 is plotted on the x-axis and associated pLDDT value per residue for each model is plotted on the y-axis. Panel A shows the positively charged residues changed to either a negatively charged residue or uncharged residue and its effect on model confidence. Panel B shows negatively charged residues changed to either positive or uncharged residues and resulting confidence values per residue. Panel C shows charged to uncharged variants. Panel D shows uncharged to positively and negatively charged variants. For all panels, the WT (black) is also plotted for comparison in model confidence. A lower confidence value compared to the WT indicates structural uncertainty and disorder promoting effects of the variant. Confidence cutoff is indicated by a dashed gray line. Models with values below this threshold are significantly destabilized by the resulting variant. Stabilizing variants have pLDDT values that lie above those of the wild-type.

**Table 2.** Query sequences for charged residues of long C-terminal domain and Helix 8.

| Sequence change | hA2a Region | Substitution |
| --- | --- | --- |
| uncharged (L, I, T, F, Y, A, Q, S, V) → negatively charged | Helix 8 | Aspartate (D) |
|  |  | Glutamate (E) |
| uncharged (L, I, T, F, Y, A, Q, S, V) → positively charged | Helix 8 | Lysine (K) |
|  |  | Histidine (H) |
| Pos (H, K, R) → negatively charged | Helix 8 | Aspartate (D) |
|  |  | Glutamate (E) |
| Neg (D, E) → positively charged | Helix 8 | Lysine (K) |
|  |  | Histidine (H) |
| Neg (D, E) → uncharged | Helix 8 | Leucine (L) |
|  |  | Glycine (G) |
| Pos (H, K, R) → uncharged | Helix 8 | Leucine (L) |
|  |  | Glycine (G) |
| charged → uncharged | Helix 8 | Leucine (L) |
|  |  | Glycine (G) |
| C-terminal disordered region :<br>(289-412) - long<br>Helix 8 | <sup>1(312)</sup>EPFKAAGTSARVLA AHGSDGEQVSLRLNGHPPGVWANGS<br>APHPERRPNGYALGLVSGGSAQESQGNTGLPDVELLSHELKGVC<br>PEPPGLDDPLAQDGAGVS<sup>101(412)</sup><br><sup>1(289)</sup>AYRIREFRQTFRKIIRSHVLRQQ<sup>23(311)</sup> |  |

\*confidence values

In general, fewer destabilizing effects are seen when negatively charged residues are substituted with positively charged or uncharged residues (Fig. 6 – Panel B). Leucine substitutions also promote helical structure when substituted for charged residues compared to the WT (Fig. 6 – Panel C). Glycine has the opposite effect, disrupting helical structure and reducing certainty in the model. Again, the resulting effects of these respective substitutions are consistent with benchmark findings for the poly-AA predictions. Most of the changes in model confidence due to sequence variations and mutations lie outside the margin of error for each prediction and are, therefore, significant.

## CONCLUSIONS

This analysis focuses on substitutions using both structured and disordered domains of the human adenosine A2a receptor, a GPCR important for cell signaling and central to many physiological processes. These well-characterized domains enable us to probe the contribution of each residue to the structure of the protein, to provide insight into the fundamental principles that govern protein disorder. This information provides a steppingstone from which we can continue our attempts to understand the underlying factors that drive protein aggregation in disordered sequences and ultimately what leads to irreversible aggregation seen in some of the most debilitating diseases. While we can draw conjectures about how selected substitutions “alter” or destabilize the predicted structures of the disordered C-terminal domain and the 8^th^ C-terminal helix by introducing uncertainty into the models, ColabFold-AF2 is not able to determine changes in thermodynamic parameters governing any stabilizing or destabilizing forces. Nonetheless, many of the insights generated can be reconciled with previous studies lending support to the reported findings here. This study paves the way for analyses probing additional factors that impact polypeptide disorder to further uncover intramolecular and cellular conditions that lead to their irreversible aggregation.

The disordered structures depicted here, which were generated in ColabFold, do not represent absolute static structures, nor do they represent the true conformation of these polypeptide sequences. They do, however, provide important insights about how residues in protein primary structure contribute to a preferred disordered structural state, and we have begun to unravel how residues work in concert to maintain their structured or unstructured characteristics. Using this analysis, we can also draw insights about fundamental factors that contribute to intrinsic disorder. Further, an analysis investigating protein-protein interactions using programs like AlphaFold-Multimer or even the most recently updated AlphaFold 3, might help to elucidate additional properties that facilitate the intermolecular interactions that lead to the aggregation events that trigger uncontrolled and irreversible high-order oligomer formation^43^.

These findings also do not take into account changes in disordered structure compaction and chain elongation, biophysical properties of IDPRs that are beyond the scope of ColabFold ^44–47^. While IDPRs generally assume elongated, dynamic structures compared to their structured counterparts, variations in special compaction and chain length as a function of amino acid composition are an important consideration when predicting structural conformation of IDPRs and perhaps even their propensity to aggregate. Protein modifications such as posttranslational modifications (PTMs) might also play a role in stabilizing structural disorder; however, these features are not captured here ^48^.

### Summary of order and disorder-promoting residues

The order and disorder-promoting tendency of each amino acid was tested by predicting the structure of a 25-mer polypeptide for each residue. The predictions are consistent with the structure-induced effects observed when residues were then substituted into either the disordered C-terminal domain or structured helix 8 of a well-characterized protein, hA2a.

#### Proline is a disorder-promoting helix disruptor

From the collective data, we conclude that the extent of helical disruption induced by proline can be detected by the ColabFold program, but it depends on the surrounding sequence residues in the α-helical structure. It may be more difficult to detect point mutations in larger, more complex protein structures where numerous intramolecular interactions contribute to the final structure prediction. To circumvent this, it may be helpful to simply analyze a specific region of interest where mutations either occur naturally or where they can be introduced to evaluate the structural contributions of a single residue.

#### Leucine and alanine are order-inducing helix promoters

They induce localized changes to the regions of substitution but seem to have little effect on the global structure where it remains overwhelmingly disordered. A closer systematic analysis may help uncover additional factors that contribute to the disordered state. For example, a more extensive analysis of charge distribution ^49–51^. Having charged or uncharged residues present, does not seem to dictate whether a polypeptide adopts a structured or unstructured conformation. Clearly, there are other factors at play. The surrounding residues in the polypeptide chain matter: they can help stabilize secondary structure and overcome the effects of disorder-promoting residues. **I**n the future, we aim to probe further into the details of the rules governing the interplay between flanking/surrounding residues and sites of substitution for helix disrupting residues. This will necessitate an extensive biophysical analysis of each residue along with careful consideration of their thermodynamic contributions to overall stabilizing interactions of secondary structure. Lastly, we have not considered the impact of side-chain steric effects. This is an important consideration as well. Aspartate is highly disruptive to helical structure, while other charged residues; glutamate, lysine and arginine, are not (Supplementary Table S1). A simple physicochemical analysis of their properties is insufficient to fully explain each individual contribution to structure.

ColabFold has enabled us to uncover behavioral predictions of order- and disorder-promoting amino acids in a single polypeptide chain and their induced effects on secondary structure. However, challenges remain; extracting meaningful information from predictions that already carry substantial uncertainty as in the case of IDPRs. While we have attempted here to make predictions about disorder from primary sequence information, these predictions cannot tell us one way or the other about the disease pathology of the query sequences tested or their propensity to aggregate. The findings in this study recapitulate the need for experimental validation. Experimental validation will strengthen the confidence with which we use structure prediction programs like ColabFold-AF2 and other iterations.

## Abbreviations

IDP/Rs: intrinsically disordered proteins/regions
A_2A,_ hA2a: human adenosine
WT: wild-type
AF2: AlphaFold2
GPCRs: G-protein coupled receptors
cryo-EM: cryo-electron microscopy
NMR: nuclear magnetic spectroscopy

## ACKNOWLEDGEMENTS

The author acknowledges the use of ChatGPT (version 3.5) by OpenAI (https://chatgpt.com/) to generate a draft of select sections of the Introduction. The author is credited with generating the appropriate ideas, prompt inputs, critical evaluation and referencing and citing previous work. All content in the finalized manuscript was edited for clarity, scientific accuracy and stylistic prose by the author. ChatGPT was never used to generate ideas, analyze and interpret data or draw any conclusions using the results of this study.

## Supplementary Information

**Table S1.**
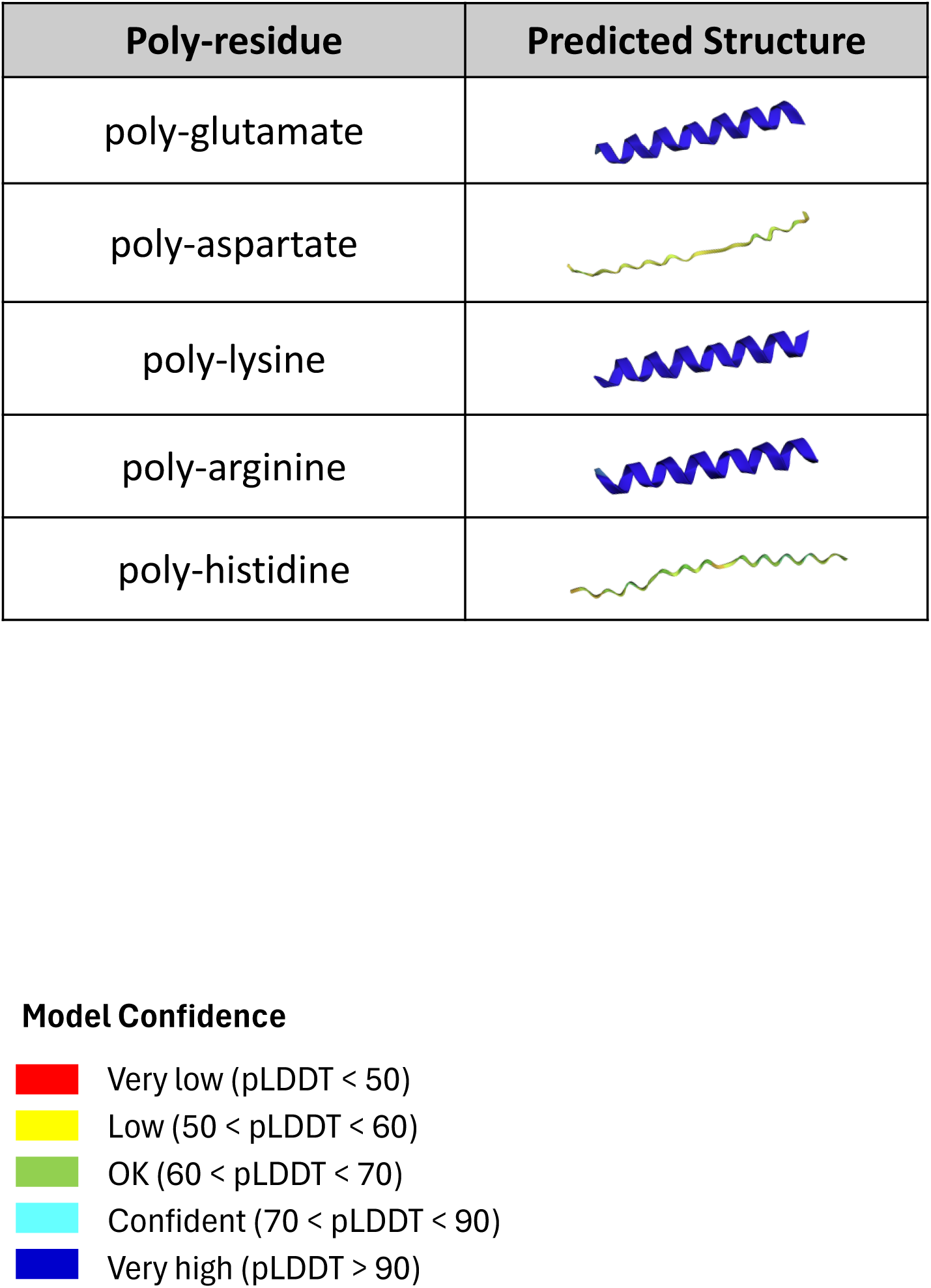
Charged poly-AA peptides.

| Poly-residue | Predicted Structure |
| --- | --- |
| poly-glutamate |  |
| poly-aspartate |  |
| poly-lysine |  |
| poly-arginine |  |
| poly-histidine |  |

**Table S2.**
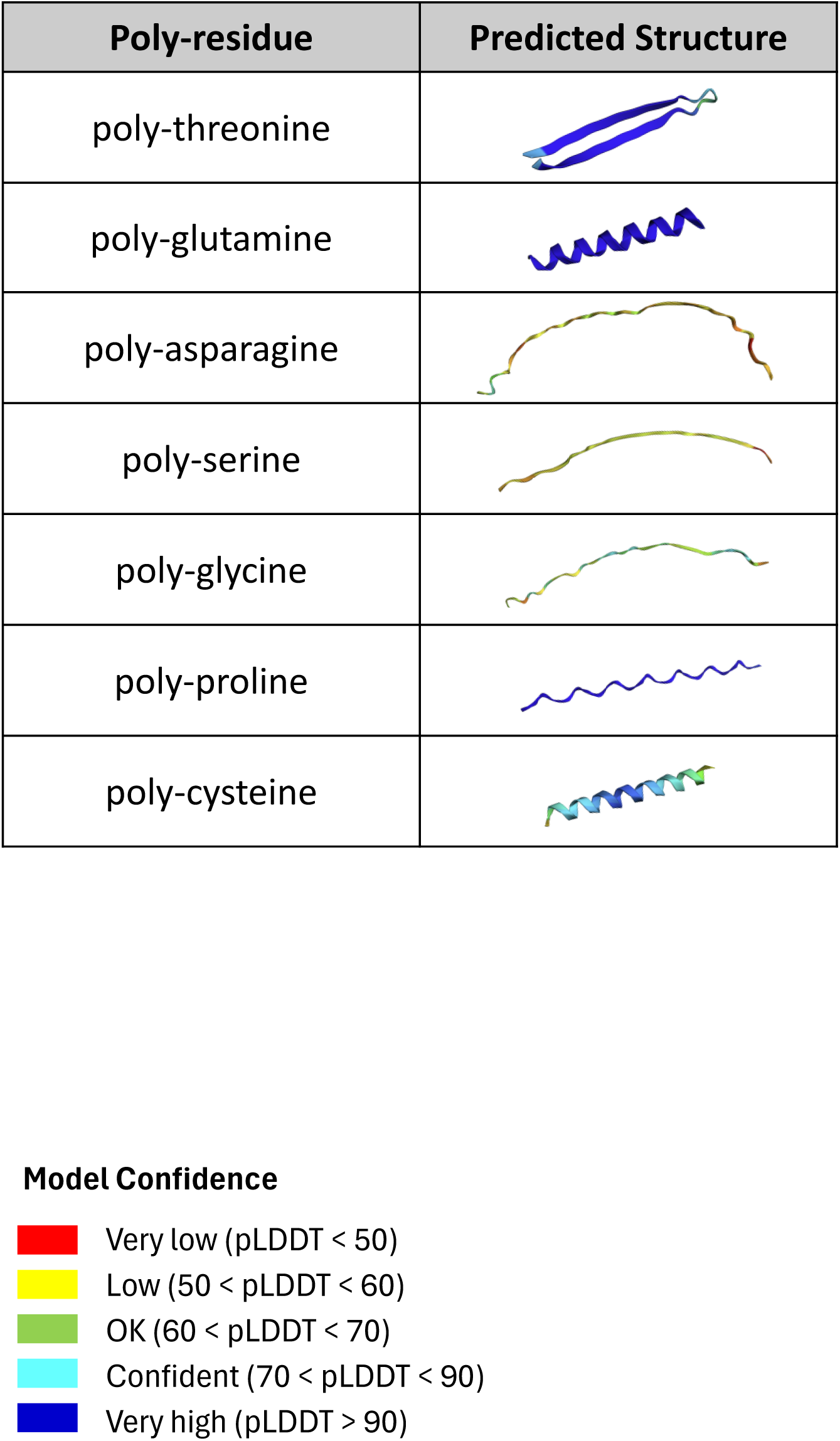
Polar poly-AA peptides.

| Poly-residue | Predicted Structure |
| --- | --- |
| poly-threonine |  |
| poly-glutamine |  |
| poly-asparagine |  |
| poly-serine |  |
| poly-glycine |  |
| poly-proline |  |
| poly-cysteine |  |

**Table S3.**
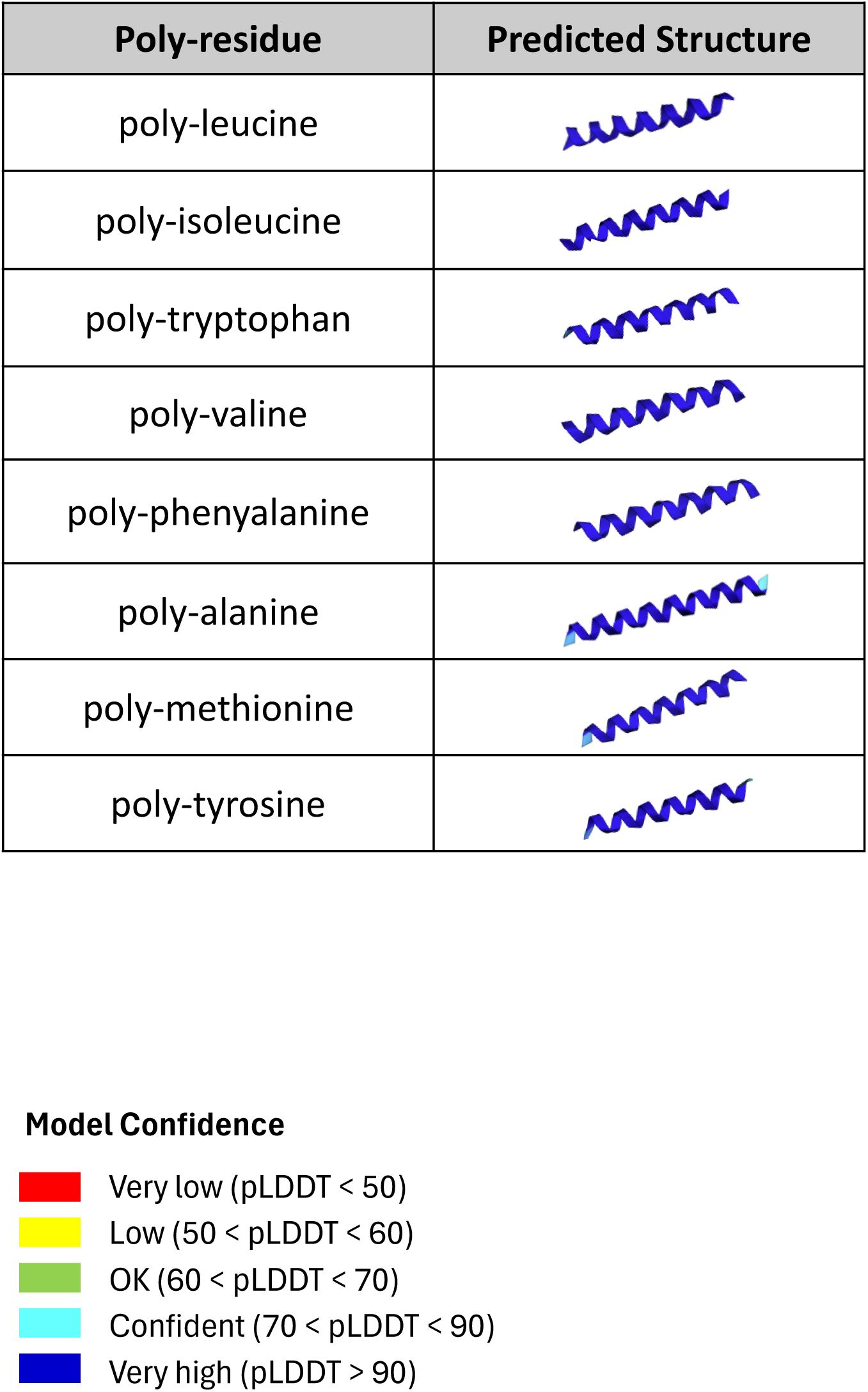
Nonpolar poly-AA peptides.

**Figure S1.**
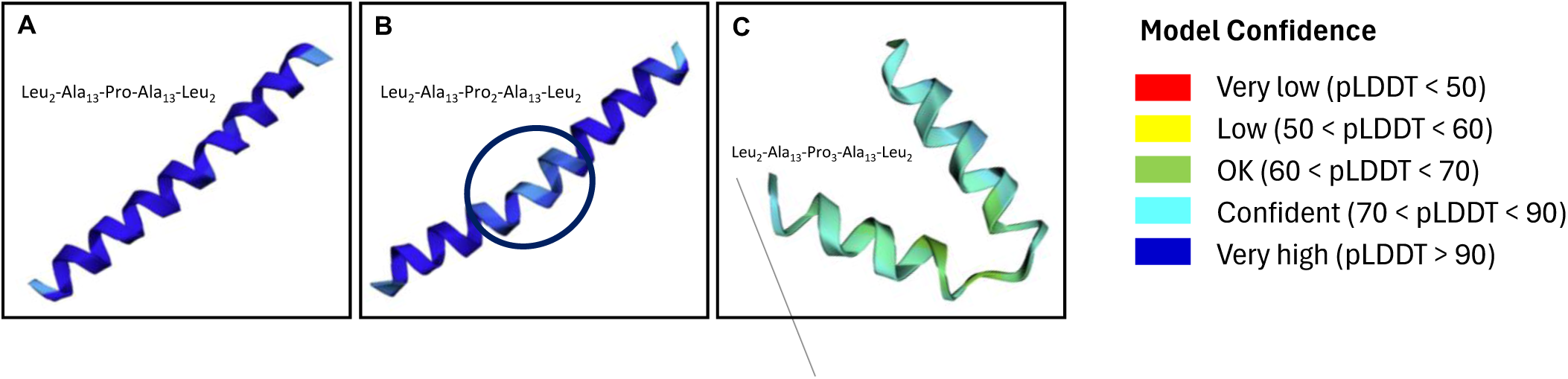
Effects of proline on secondary structure predicted with ColabFold. A) Polyleucine/polyalanine hybrid helix with one proline residue introduced, B) two consecutive proline residues introduced results in a slight kink to the helix structure (circled), C) three consecutive proline residues results in a near complete turn within the helix.

**Figure S2.**
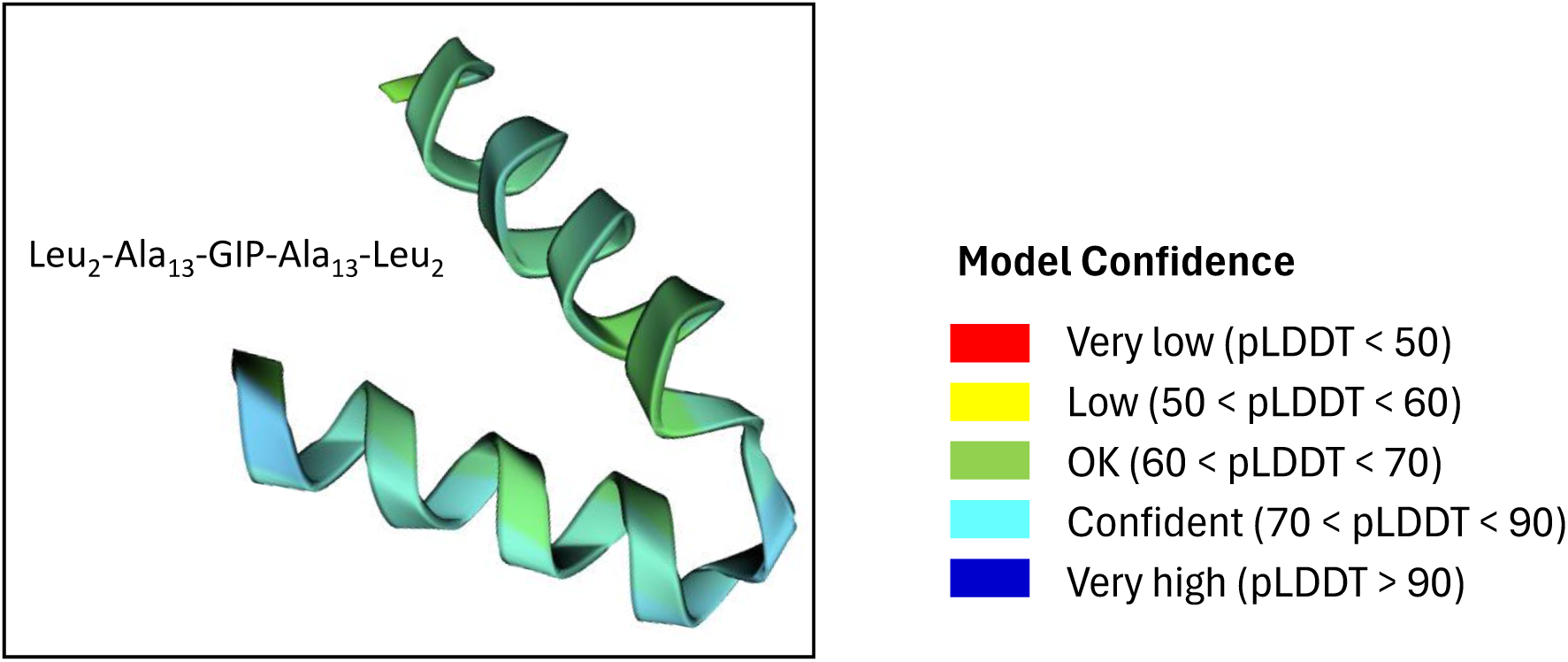
Predicted structure of three-residue (G I P) turn introduced to a polyleucine/polyalanine hybrid helix. The three-residue turn includes glycine and proline, two helix disrupting residues.

**Figure S3.**
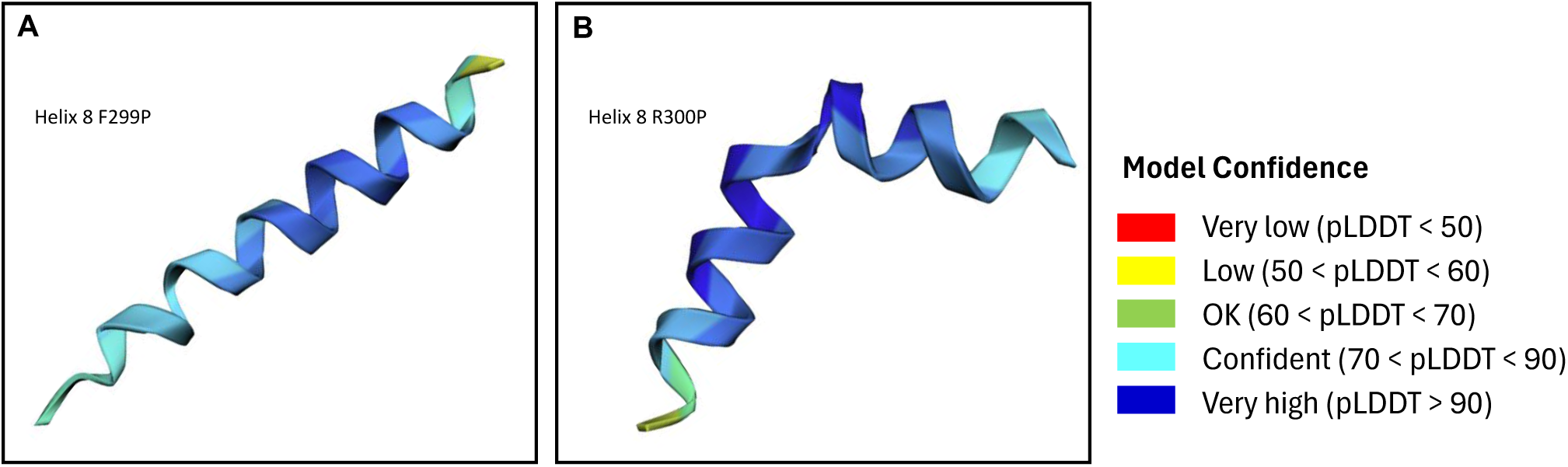
Single proline substitution at two adjacent positions in Helix 8 of hA2a. A) position F299 B) position R300. The position of the substitution and flanking residues influence the predicted structure. Structures are shown N terminus to C terminus.

**Figure S4.**
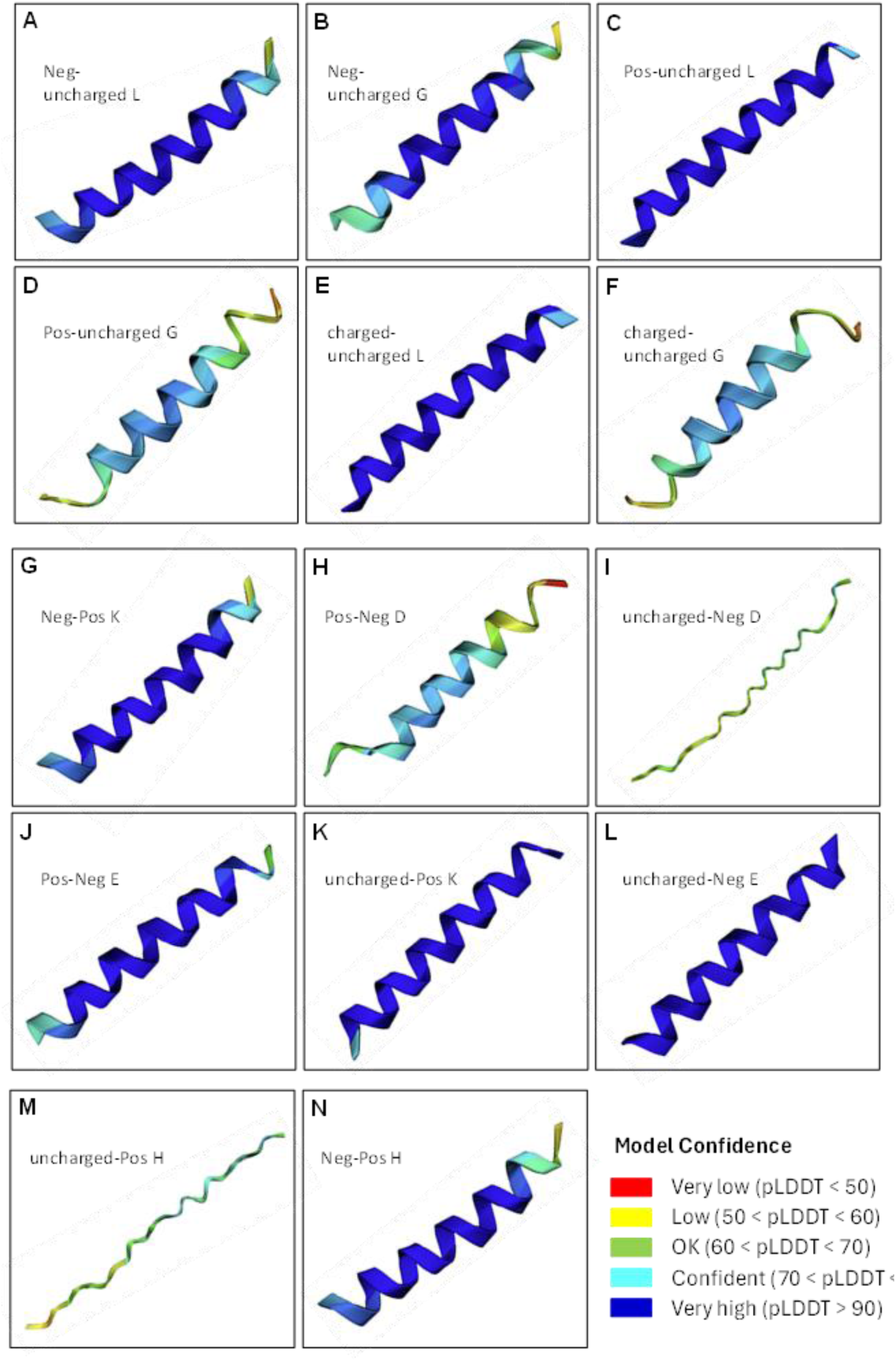
ColabFold predictions of charge-substituted variants in Helix 8. The model structures in each panel are shown N terminus to C terminus from left to right. Aspartate is most disruptive to helical structure and considered disorder-promoting, panels H & I. Glycine is also disorder-promoting, panels D & F. Leucine, glutamate, lysine and are order-promoting, panels A, C, E, G, J, K & L. Histidine can be considered disorder-promoting and the extent of disruption depends on the extent of substitution, where heavy substitution (5 residues or more) leads to near complete disruption of helix 8, panel M. While a single substitution only slightly disrupts the helical structure, panel N. Structures are shown from N terminus to C terminus.

